# Differential expression and localization of thermosensitive Transient Receptor Potential Vanilloid (TRPV) channels in the mature sperm of white pekin duck (*Anas platyrhynchos*)

**DOI:** 10.1101/2020.02.10.941732

**Authors:** Rakesh Kumar Majhi, Ashutosh Kumar, Sunil C. Giri, Chandan Goswami

## Abstract

Sperm cells have the ability of precise chemotactic and thermotactic movement which is crucial for fertilization, yet the key molecules involved in the detection of different chemical and physical stimuli and guide the sperm cells for proper navigation are not known. This aspect is more complex as each species have their own reproductive identity. Never-the-less, Ca^2+^-signaling and thus a series of Ca^2+^-channels seem to coordinate in order to regulate different functions mediated by sperm cells. However, such aspects are controlled by different Ca^2+^ channels and have species-specific differences. In this work we explored if TRPV channels are endogenously expressed in the mature spermatozoa obtained from avian species. We have used the sperm cells of white pekin duck (*Anas platyrhynchos*) as a representative avian species to explore the endogenous expression and localization of different TRPV channels. Western blot analysis (WB), flow cytometry, confocal imaging and super resolution imaging was performed for the characterization. Our results strongly suggest the expression and distinct localization of different TRPV channels in the sperm cells. All these TRPV channels are mainly absent in the head region. Only TRPV3 and TRPV4 are sparsely present in the neck region enriched with mitochondria. All these channels (TRPV1-6) are present in the tail region. The differential localization of TRPVs in duck sperm indicate their respective functions relevant in fertilization process of avian sperm. These findings may also have commercial importance in poultry production, cryopreservation of sperm as well as conservation of endangered species through artificial insemination.

## 1. Introduction

Most of the mature and healthy sperm have the ability to swim a long distance within the female reproductive tract in order to successfully fertilize the egg. During this journey, sperm is guided through chemotaxis, rheotaxis and thermotaxis [1–2]. The capacity to detect and respond to temperature changes by sperm is a critical determinant of species evolution too [3]. Sperm cells are believed to be highly thermosensitive in nature. In fact human sperm have been shown to respond to even <0.0006 °C temperature difference [4]. Practical utility of thermotaxis ability has been demonstrated recently, where capacitated murine and human spermatozoa selected through swim up procedure across a temperature gradient of 35°C to 38°C were observed to show better fertilizing ability [5]. This precise thermosensitivity also accords well with the existence of temperature gradient between oviduct isthmus and ampulla [2].

Timing of each of these responses is critical and premature or inappropriate activation of these events can lead to failure in fertilization. Since sperm cells are mostly transcriptionally and translationally inactive, all cellular activities within it are carried out by the pool of proteins that are generally inherited during differentiation of spermatozoa and these proteins are responsible to regulate sperm functions via secondary messengers like intracellular Ca^2+^ [6–8]. In ejaculated sperm cells, intracellular Ca^2+^-influences motility, chemotaxis, capacitation, hyperactivation and acrosome reaction [9–13]. The Ca^2+^-signaling is initiated and maintained by Ca^2+^-permeable ion channels present in the sperm. Besides regulating calcium signaling, these channels also sense the chemical cues and thermal environment of the sperm and modulate the sperm in accordance with the stimuli.

The ability to respond minute differences in temperature suggests that mature sperms are equipped with the specialized thermo-responsive machineries at the molecular level. Such molecules can detect specific temperature, induce proper cell signaling leading to sperm movement in order to undergo thermotaxis. Thermosensitive Transient Receptor Potential (TRP) channels are likely to be present in mature sperm of different species and these channels are important for all these process. TRP channels are the most diversified group of ion channels and is generally characterized as non-selective to cations, implying that they can conduct influx of several monovalent and divalent cations upon activation, including Ca^2+^ ions [14]. TRP channels are characterized by the presence of six transmembrane helices, an intracellular N-terminus and a C-terminus tail. The cation conducting pore loop is present between the 5^th^ and 6^th^ transmembrane helices. TRP channels are classified into 7 subfamilies based on their amino acid sequence and structural homology [15]. The TRPVs (‘vanilloid’) have been named after the founding member Vanilloid Receptor 1 (TRPV1) and consists of 6 members TRPV1, TRPV2, TRPV3, TRPV4, TRPV5 and TRPV6. While TRPV1, TRPV2, TRPV3, TRPV4 are non-selective for cations and are thermosensitive in nature, TRPV5 and TRPV6 are relatively more selective for Ca^2+^ ion and not thermosensitive in nature. Efforts by various groups has led to increasing knowledge about their localization and functional relevance of specific TRP channels in various cell types, including sperm cells.

Indeed, heat sensitive TRPV1 has been shown to be functional in fish, mice, boar, bull and human sperm [16–20]. TRPV1 has been shown to be important for progesterone-induced sperm oocyte fusion and TRPV1 activator anandamide can capacitate bull sperm [19, 21]. TRPV1 has also been demonstrated to mediate thermotaxis of human sperm [20]. Interestingly TRPV4 has been also shown to mediate thermotaxis in mouse sperm and initiate the initial depolarization event for capacitation in human sperm [22–23]. TRPV4 has been reported by us to be present in all vertebrate sperm [24]. TRPV2, TRPV3, TRPV5 and TRPV6 haven’t been reported in the sperm of any species so far. In spite of several reports, TRP channels, including TRPV channels have remained relatively unexplored in avian sperm. Previously we have described the unique morphology of mature duck sperm and presence of different modified tubulin in such cells [25]. Since endogenous expression and localization of these channels can be correlated to sperm function, therefore, in this work we explored the presence of TRPV channels in avian (duck sperm). We noted distinct pattern of localization of the thermosensitive TRPV1, TRPV2, TRPV3, TRPV4 channels and highly calcium-selective TRPV5, TRPV6 channels along the duck sperm.

## 2. Materials and Methods

### 2.1 Collection of mature spermatozoa from duck

Sperm was collected as described previously [25–26]. Briefly, white pekin ducks (*Anas platyrhynchos*) maintained in the duck house (Central Avian Research Institute, Bhubaneswar, India) were massaged manually for semen collection by trained professionals in accordance with the guidelines of the Institutional animal ethics committee. No animals were harmed or sacrificed for this work. Mature sperm were collected into clean sterile collection vials and either fixed with 4% Paraformaldehyde (PFA) for immunostaining or were incubated at 37°C immediately after ejaculation staining with Mitotracker-Red dye to visualize the neck region. Ejaculated sperm were immediately processed for Western blotting. All microscopic, flow-cytometric and biochemical analysis were performed at the NISER, Bhubaneswar. All experiments were done according to the approval of the animal ethics committee of NISER (NISER-IAEC/SBS-AH/07/13/10).

### 2.2 Mitotracker red labelling

Freshly ejaculated duck sperm were incubated for 20 min with mitochondrial potential dye Mitotracker-Red (5 μM; Life Technologies) at 37 °C and then fixed with 1:1 ratio (v/v) of 4% PFA for 20 min at room temperature. Mitotracker-Red stained samples were washed thrice with Phosphate Buffered Saline (PBS, pH 7.4, Himedia Labs) and the nucleus was counterstained with DNA labelling dye 4’,6-diamidino-2-phenylindole (DAPI, 5μg/ml) for 15 minutes. The mitochondrial and head regions of sperm were stained to enable proper characterization of the localization pattern of TRPV channels. Differential Interference Contrast (DIC) images further help in this process by enabling visualization of the entire tip-of-head to end-of-tail region as shown by us previously [25].

### 2.3 Antibody staining and imaging

Sperm were processed for immunofluorescence microscopy analysis as described earlier [25]. For immunostaining, the fixed sperm were washed with PBS thrice by centrifugation at 800 g for 5 minutes each time. The sperm were then permeabilized with 0.1% TritonX-100 (Sigma-Aldrich) in PBS for 5 minutes, followed by blocking with 5% Bovine Serum Albumin (BSA, Sigma Aldrich) for 1 hour. All above procedures were performed at room temperature (25°C). Sperm were then incubated overnight with primary antibodies against TRPV channels dissolved in 2.5% BSA in PBS at 1:200 dilution. Primary antibodies against TRPV channels were Anti-TRPV1 antibody (Ab1: Alomone Labs, Cat. No. ACC-030; Ab2: Sigma Aldrich, Cat. No. WH0007442M1); Anti-TRPV2 antibody (Ab1: Alomone Labs, Cat. No. ACC-032; Ab2: Calbiochem, Cat. No. PC421); Anti-TRPV3 antibody (Ab1: Alomone Labs, Cat. No. ACC-033; Ab2: Sigma Aldrich, Cat. No. SAB-1300540; Ab3: Sigma Aldrich, Cat. No. SAB1300539); Anti-TRPV4 antibody (Ab1: Alomone Labs, Cat. No. ACC-034; Ab2: Sigma Aldrich, Cat. No. T9075; Ab3: Sigma Aldrich, Cat. No. HPA 007150); Anti-TRPV5 antibody (Ab1: Alomone Labs, Cat. No. ACC-035; Ab2: Sigma Aldrich, Cat. No. T3076); Anti-TRPV6 antibody (Ab1: Alomone Labs, Cat. No. ACC-036). Validation of antibodies was done through peptide blocking experiments, where possible. For antibody blocking experiments, the antibodies were pre-incubated with their respective antigenic peptides (Alomone Labs) for 30 minutes at 1:1 (w/w) ratio, prior to adding to the sperm solution. The blocking peptides (all from Alomone labs) had the following sequences: TRPV1 (EDAEVFKDSMVPGEK); TRPV2 (KKNPTSKPGKNSASEE); TRPV4 (CDGHQQGYAPKWRAEDAPL); TRPV6 (NRGLEDGESWEYQI). After incubation, the sperm were washed thrice with PBS-T (0.1% Tween20 in PBS) thrice by centrifugation. Alexa Fluor-488 labeled anti-mouse or anti-rabbit antibodies (Molecular probes) were used as secondary antibodies at 1:1000 dilution and incubated for 1hour. The nucleus were counterstained with DAPI (4’,6-diamidino-2-phenylindole, 5μg/ml) for 15 minutes. After washing thrice with PBS-T and dissolving in FACS buffer (PBS with 0.01% sodium azide and 0.1% BSA) the cells are ready for imaging and flow cytometry. All images were acquired on confocal laser-scanning microscope (LSM-780, Zeiss) using a 63X-objective and analyzed with the Zeiss LSM image examiner software as described previously [16]. The super resolution images of duck sperm were acquired using super resolution structured illumination microscopy (SR-SIM; Zeiss ELYRA PS.1, (Carl Zeiss) and processed using Zen 2 Blue software (Carl Zeiss).

### 2.4 Flow cytometry

Flow cytometry analysis was performed using BD FACS Calibur flow cytometer (BD Biosciences) equipped with two lasers emitting at 488 and 635 nm and as described previously [24]. Forward scatter (FSC) signal was collected by a photodiode, while Side scatter (SSC) was detected by a PMT with Brewster-angle beam splitter, both equipped with 488/10 bandpass filter. The fluorescent signal (only green fluorescence emitted by AlexaFluor 488 used to detect TRPV channels) were collected by a fluorescence collection lens FL1: excitation 488 nm, emission 530nm (530/30 nm bandpass filter). Sample density at time of acquisition was 0.5 x 10^5^ cells/ml and 10,000 sperm were acquired in live-gated population (P1) per sample at medium sample flow rate. The instrument was calibrated with BD Calibrate beads routinely as specified by the manufacturer. Unstained samples were utilized to set up -ve events compensations, and to select regions of interest, while multi-color fluorescence compensations were not required since the samples were single-stained with only Alexa Fluor 488 (detected in FL1 channel). The data was analyzed using Cell Quest Pro Software (BD Biosciences). Dot plot values reflected the percentage of sperm that express TRPV channels. Mean Fluorescence Intensity (MFI) was analyzed as a numerical value (in arbitrary unit) to comment on the relative expression level of TRPV channel in each sample population.

### 2.5 SDS PAGE and Western Blot analysis

Western blotting was performed as described by us previously, but with minor modifications [16, 24]. Freshly collected duck sperm was diluted in 1XPBS and centrifuged at 800 g for 5 minutes in 25°C. The sperm pellet was diluted in 1× PBS supplemented with 2X Protease inhibitor cocktail (Sigma Aldrich) and 5X Laemmli gel sample buffer was added to it. The samples were boiled at 95°C for 5 minutes and subsequently analyzed by 10% sodium dodecyl sulfate PAGE (SDS-PAGE) according to Laemmli [27]. The proteins were transferred to PVDF membrane by semi dry transfer apparatus in 45 minutes (Millipore). The blots were blocked for 1 hour in 5% (w/v) dry skim milk dissolved in TBST (20 mM Tris [pH 7.4], 0.9% (w/v) NaCl, and 0.1% (v/v) Tween 20). Then the membranes were incubated with antibodies against TRPV channels at 1:200 dilution in cold room for overnight. After washing thrice in TBST, the membranes were incubated with horse-radish peroxidase-conjugated secondary antibody raised against rabbit or mouse (GE Healthcare) for 1 h at room temperature (25°C). For antibody blocking experiments, the antibodies were pre-incubated with their respective antigenic peptides (Alomone Labs) for 30 minutes at 1:1 (w/w) ratio, prior to adding to the membranes. The membranes were again extensively washed in TBST and bands were visualized using ChemiDoc XRS+ system (Bio-Rad) by using enhanced chemiluminescence according to the manufacturer’s instructions (Super-Signal West Femto Maximum Sensitivity Substrate, Thermo Scientific).

### 2.6. Statistical analysis

All flow cytometry experiments were performed in triplicates. Statistical analyses were performed by using GraphPad Prism 5 software. Significant differences were determined between unstained and TRP channel stained samples by performing unpaired Student’s T-test after confirming normal distribution and homogeneity of variances of the data. The results are expressed as mean ± standard error of mean (SEM). Data was considered to be significant at *P* < 0.05.

## 3. Results

### 3.1 TRPV channels are endogenously expressed in duck sperm

To detect the endogenous expression of TRPV channels in duck sperm, we performed western blotting analysis using duck sperm extract. Wherever possible, we have used antigenic peptide for the antibodies used to show their specificity for the TRPV channel being investigated. In other cases, we have used two antibodies to detect the same channel, to establish the presence of these channels at the protein level in duck sperm. Using an antibody, directed against the C-terminus of TRPV1 (Alomone Labs), we detected a ~95kDa band and most of the signal was blocked upon pre-incubating the antibody with its antigenic peptide, confirming specificity of the antibody used (**Fig. 1A**). Western blot analysis (using Ab1, that detects TRPV2 C-terminus, Alomone Labs) revealed a ~86kDa band for TRPV2 and majority of the signal was blocked upon pre-incubating the antibody with its antigenic peptide, confirming specificity of the antibody used (**Fig. 1B**). Two prominent bands at ~75kDa and ~65kDa for TRPV3 were detected by western blot analysis (using two different antibodies raised against the C-terminus of TRPV3 (Ab1: Alomone Labs) and another antibody against N-terminus: Ab3 from Sigma Aldrich) (**Fig. 1C**). The presence of two bands for TRPV3 could be either due to alternate splicing, or post-translational modifications (such as phosphorylation and/or glycosylation) or proteolytic degradation of TRPV3 in duck sperm. Never-the-less, at least one band remain common in both antibodies detecting TRPV3 and the shift in bands in both blots could be due to alternate splicing, or post-translational modifications (such as phosphorylation and/or glycosylation) or proteolytic degradation of TRPV3 in the samples from different individuals. A ~95kDa band was obtained for TRPV4 by using an antibody raised against the C-terminus of TRPV4 (Ab1: Alomone Labs) and another antibody against N-terminus (Ab3 from Sigma Aldrich) (**Fig. 1D**). Western blot analysis using two different antibodies raised against the C-terminus of TRPV5 (Ab1: Alomone Labs and Ab2: Sigma-Aldrich) revealed a prominent band at ~85kDa and a faint band at ~100kDa (**Fig. 1E**). Western blot analysis (using Ab1: against the C-terminus of TRPV6, Alomone Labs) revealed a prominent band at ~85kDa and that could be blocked by its antigenic peptide (**Fig. 1F**). This indicated that these TRPV channels are endogenously expressed in mature sperm of duck.

**Fig. 1.**
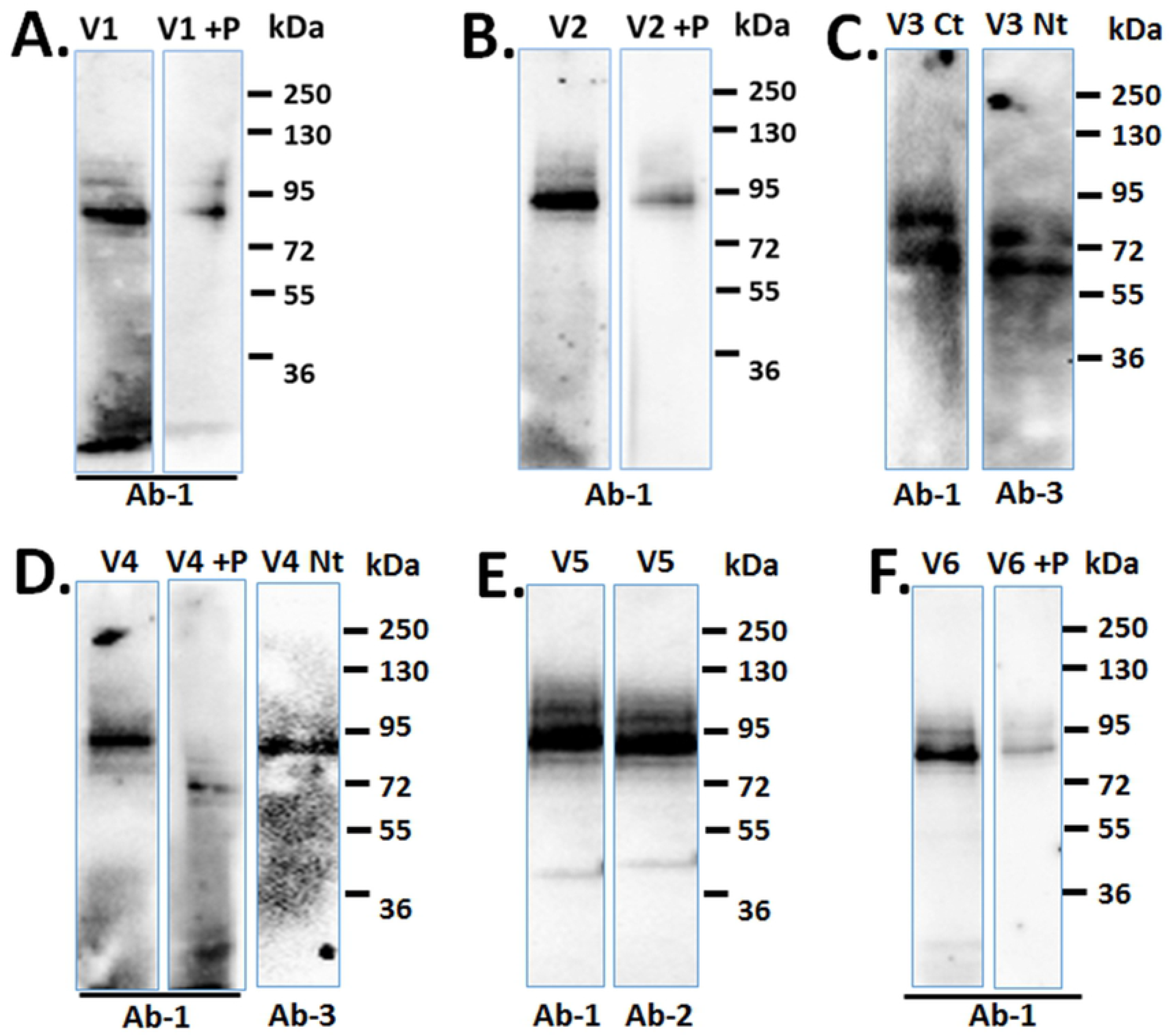
Endogenous expression of TRPV channels in duck sperm. Western blot analysis of duck sperm extracts probed with different TRPV-specific antibodies are shown. **A.** TRPV1 specific band is detected by a specific antibody (directed against the C-terminus of TRPV1, Alomone Labs) in absence but not in presence of its blocking peptide; **B.** Western blot analysis with antibody that detects TRPV2 (raised against the C-terminus, Alomone Labs). **C.** Two different antibodies detecting TRPV3 [raised against the C-terminus, (Ab1: Alomone Labs) and N-terminus (Ab3: Sigma Aldrich)] detect similar expression pattern of TRPV3. **D.** Two different antibodies raised against the TRPV4 [raised against C-terminus, Ab1: Alomone Labs) and N-terminus (Ab3: Sigma Aldrich)] detect TRPV4 at the expected size. **E.** Two different antibodies raised against the C-terminus of TRPV5 (Ab1: Alomone Labs and Ab2: Sigma-Aldrich) detects TRPV5 at expected size. **F.** A specific antibody raised against the C-terminus of TRPV6 (Ab-1: Alomone Labs) detects TRPV6 in absence but not in presence of its blocking peptide.

Notably, the difference in band size and predicted molecular weight of the TRPV channels could arise from species or individual specific differences, post-translational modifications, partial degradation of the proteins or presence of splice variants [28]. However, we have consistently obtained prominent bands near the predicted molecular weight for all channels tested. We cannot rule out the possibility of some extent of non-specific binding associated with these antibodies, which make it challenging to solely rely on western blotting. We have therefore characterized TRPV channel expression by flow cytometry and imaging, using at least two different antibodies, to gain more confidence on endogenous expression patterns.

### 3.2 Duck sperm express physiologically relevant thermosensitive TRPV channels to varying extents

Flow cytometric analysis revealed that not all sperm express all the TRPV channels to the same extent. The activation thresholds of TRPV1 (>43°C), TRPV3 (>32°C) and TRPV4 (>37°C) are within the physiological range in birds. TRPV2 (>52°C) has too high activation threshold, while TRPV5 and TRPV6 are not thermosensitive. Hence, we explored the level of TRPV1, TRPV3, TRPV4 expression levels in duck sperm using two antibodies (Ab1: from Alomone labs; Ab2: from Sigma Aldrich). Both antibodies (Ab1 and Ab2) confirmed that only around 34% of sperm express TRPV1, approx. 50% of sperm are positive for TRPV3 and nearly 80% of sperm are positive for TRPV4 (**Fig. 2**). The mean fluorescence intensity (MFI) values also show nearly 4-fold increase for TRPV1, 7-fold increase for TRPV3 and increase of approximately 8-fold for TRPV4 in comparison to unstained controls (**Fig. 2**). The MFI values are indicative of the expression level of TRPV channels being probed in duck sperm.

**Fig. 2.**
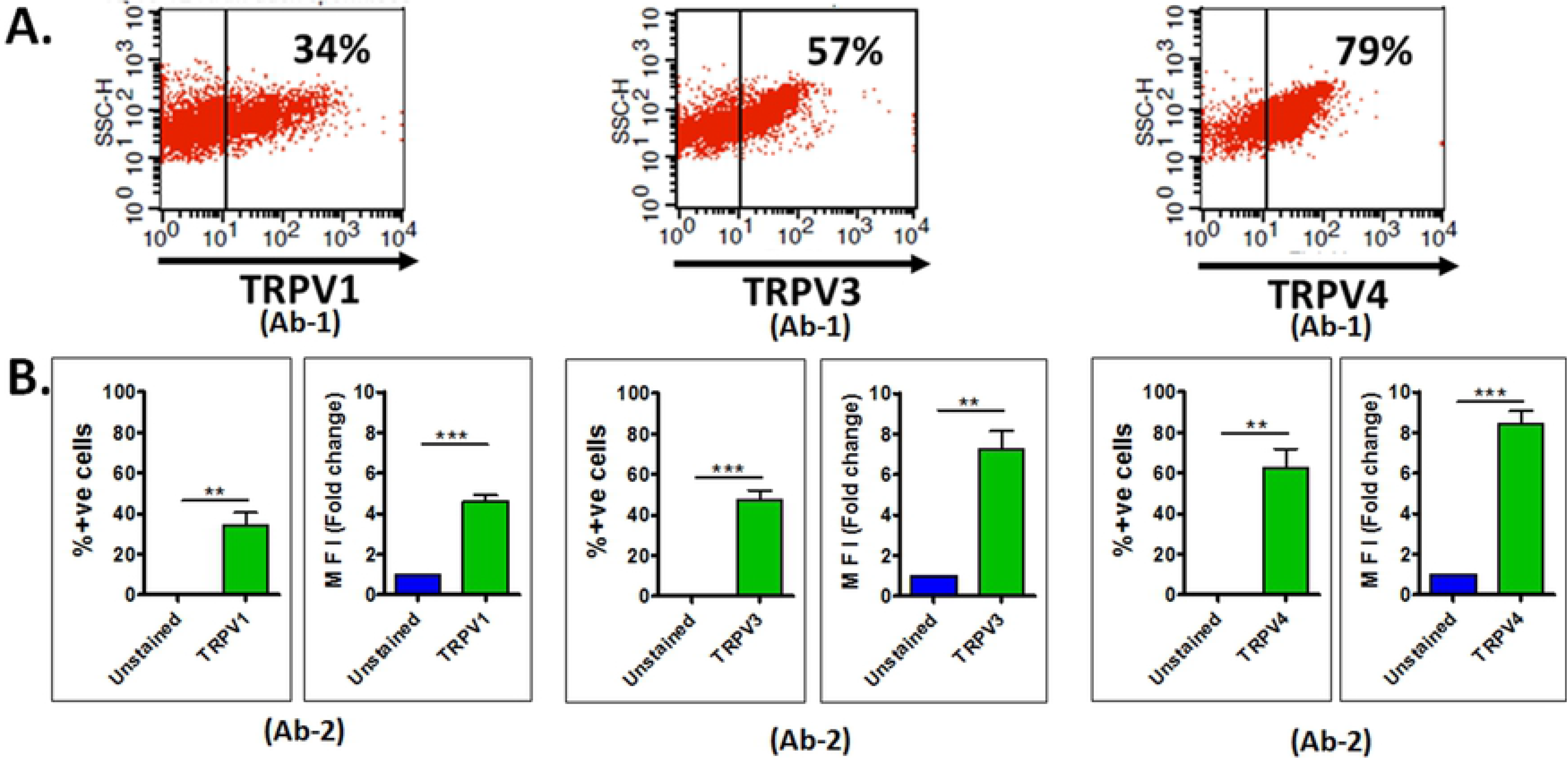
Prevalence of physiologically relevant thermosensitive TRPV channels in mature duck sperm. Flow cytometric evaluation of duck sperm stained for physiologically relevant thermosensitive TRPV channels are shown. **A.** Representative dot-plots showing percentage of cells expressing TRPV1, TRPV3, TRPV4 channels detected by Ab-1 (antibodies from Alomone labs) antibody specific for each TRPV channel. **B.** Histograms showing percentage of cells expressing TRPV channels and corresponding Mean Fluorescence Intensity (MFI) of TRPV channels detected by Ab2 antibody of each channel (from Sigma Aldrich), expressed as fold change in comparison to MFI of unstained cells. *n* = 3, unpaired T-test. ** = *P* <0.01, *** = *P* <0.001.

### 3.3 TRPV channels are differentially localized in duck sperm

To further elucidate the localization of TRPV channels in duck sperm, we have performed immunofluorescence microscopy using two different antibodies for each channel. In each of the figures (**Fig. 3–8**) we have used two different antibodies to confirm whether they reveal similar expression patterns. In the upper panels in these images, we have used DIC imaging to visualize the entire sperm, and additionally stained the mitochondrial region with MitoTracker red to demarcate the neck and used DAPI to demarcate head region to enable proper assignment of localization pattern of TRPV channels. Confocal imaging of MitoTracker red labelled duck sperm revealed presence of TRPV1 (detected by Ab1) throughout the sperm. TRPV1 is more prominent in the tail. The second TRPV1 antibody (Ab2) revealed localization of TRPV1 throughout the sperm. However, TRPV1 density was highest at the head (**Fig. 3A, B, D**). Interestingly, TRPV1 is exclusively absent in the mitochondrial region (neck region) of sperm (indicated by white arrows). SIM-based super resolution imaging (using Ab1) revealed a punctate distribution of TRPV1 throughout the sperm (**Fig. 3C**).

**Fig 3.**
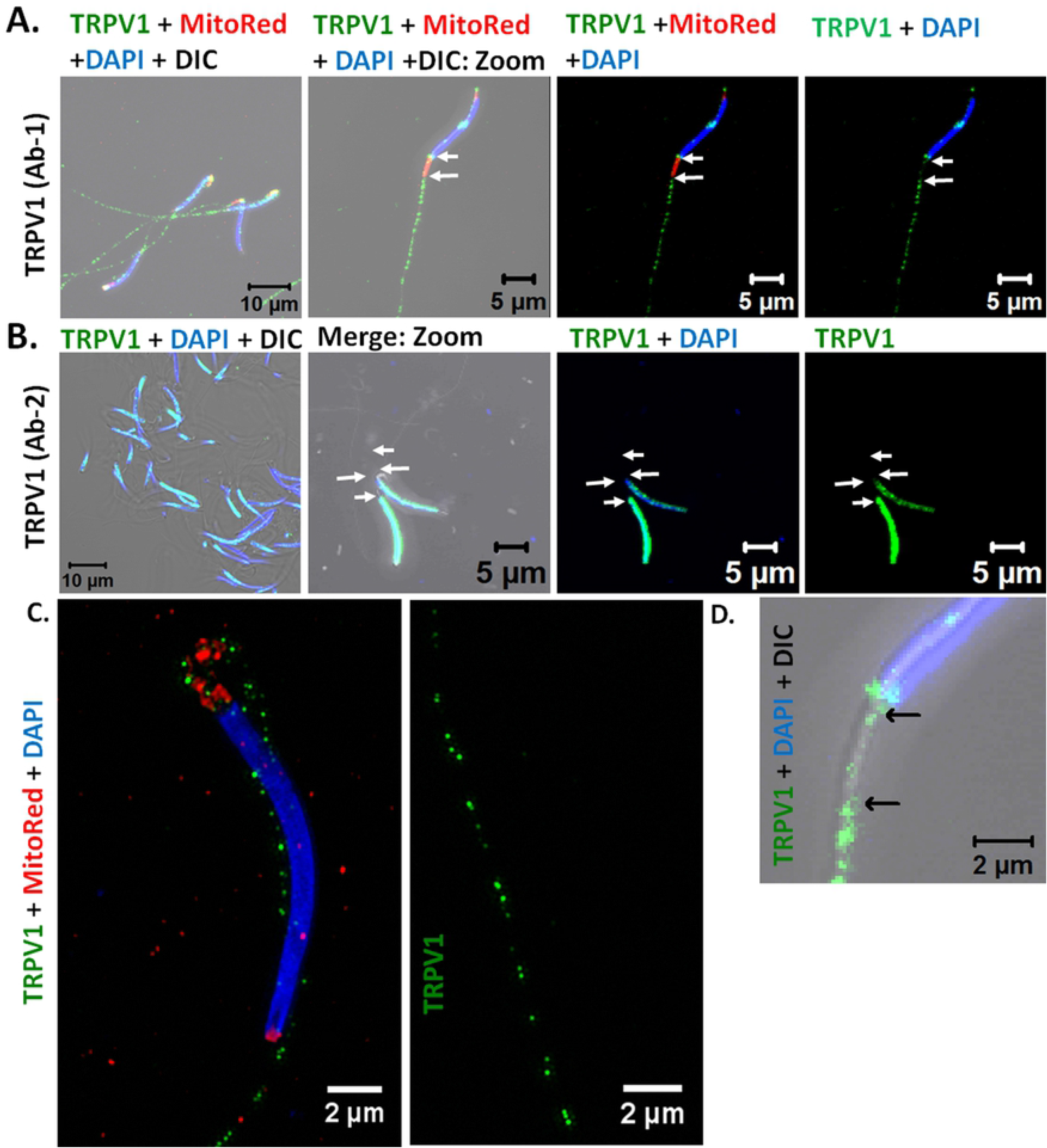
Microscopic images showing localization of TRPV1 in duck sperm. **A-B.** Confocal microscopic images depicting the localization of TRPV1 (green) as detected by two different antibodies and Nucleus (blue) by DAPI. Mitochondria (red) is labelled by Mitotracker Red dye in **A** and **C.** White arrows indicate the mitochondrial region where expression of TRPV1 is typically absent. **C**. SR-SIM images of TRPV1 localization (using Ab1 antibody) at the head (left) and tail (right) of duck sperm is shown. **D**. Zoomed up image of neck region of sperm depicting the absence of TRPV1 (green) in the neck region. The head (blue) and arrows marking the start and end point of mitochondrial region.

Using two different antibodies raised against the C-terminus of TRPV2 (Ab1 and Ab2) we found that TRPV2 is primarily present at the tail of duck sperm (**Fig. 4A, B, D**). Confocal imaging of MitoTracker red labelled duck sperm confirm the specific absence of TRPV2 in the mitochondrial region (neck region) (**Fig. 4A**).

**Fig. 4.**
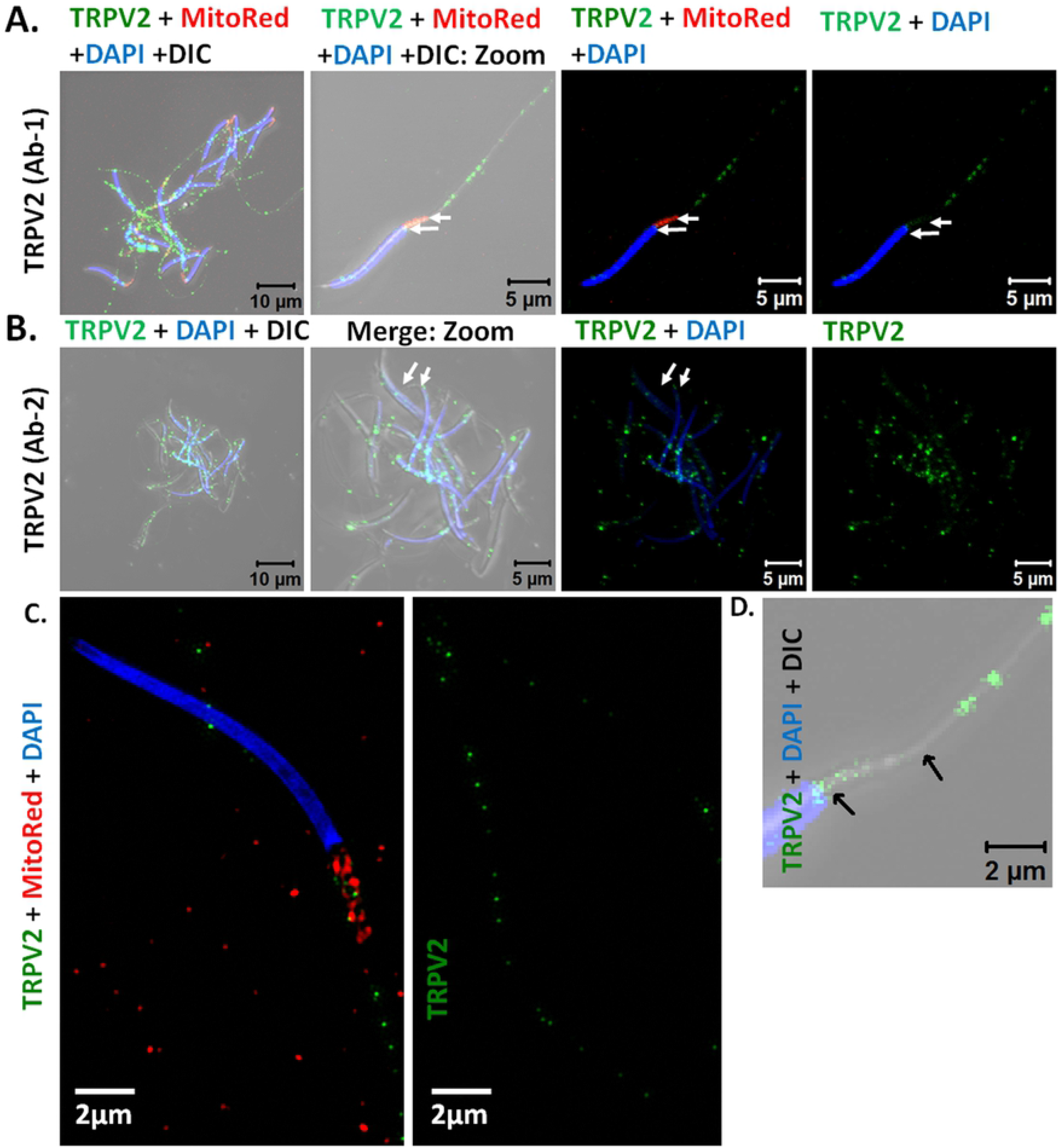
Microscopic images showing localization of TRPV2 in duck sperm. **A-B.** Confocal microscopic images depicting the localization of TRPV2 (green) as detected by two different antibodies and Nucleus (blue) by DAPI. Mitochondria (red) is labelled by Mitotracker Red dye in **A** and **C** to highlight the absence of channel expression in the mitochondrial region (indicated by arrows). **C**. SR-SIM images of TRPV2 localization (using Ab1 antibody) at the head (left) and tail (right) of duck sperm is shown. **D**. Zoomed up image of neck region of sperm depicting the absence of TRPV2 (green) in the neck region. The head (blue) and arrows mark the start and end point of mitochondrial region.

Confocal microscopy using two different TRPV3 antibodies (Ab1 and Ab2) we found that TRPV3 is primarily present at the head and tail of duck sperm (**Fig. 5A, B, D**), while very low levels of TRPV3 is present at the mitochondrial region (neck region) (**Fig. 5A**). SIM-based super resolution imaging (using Ab1) revealed a more prominent punctate distribution of TRPV3 at the tail of sperm (**Fig. 5C**).

**Fig. 5.**
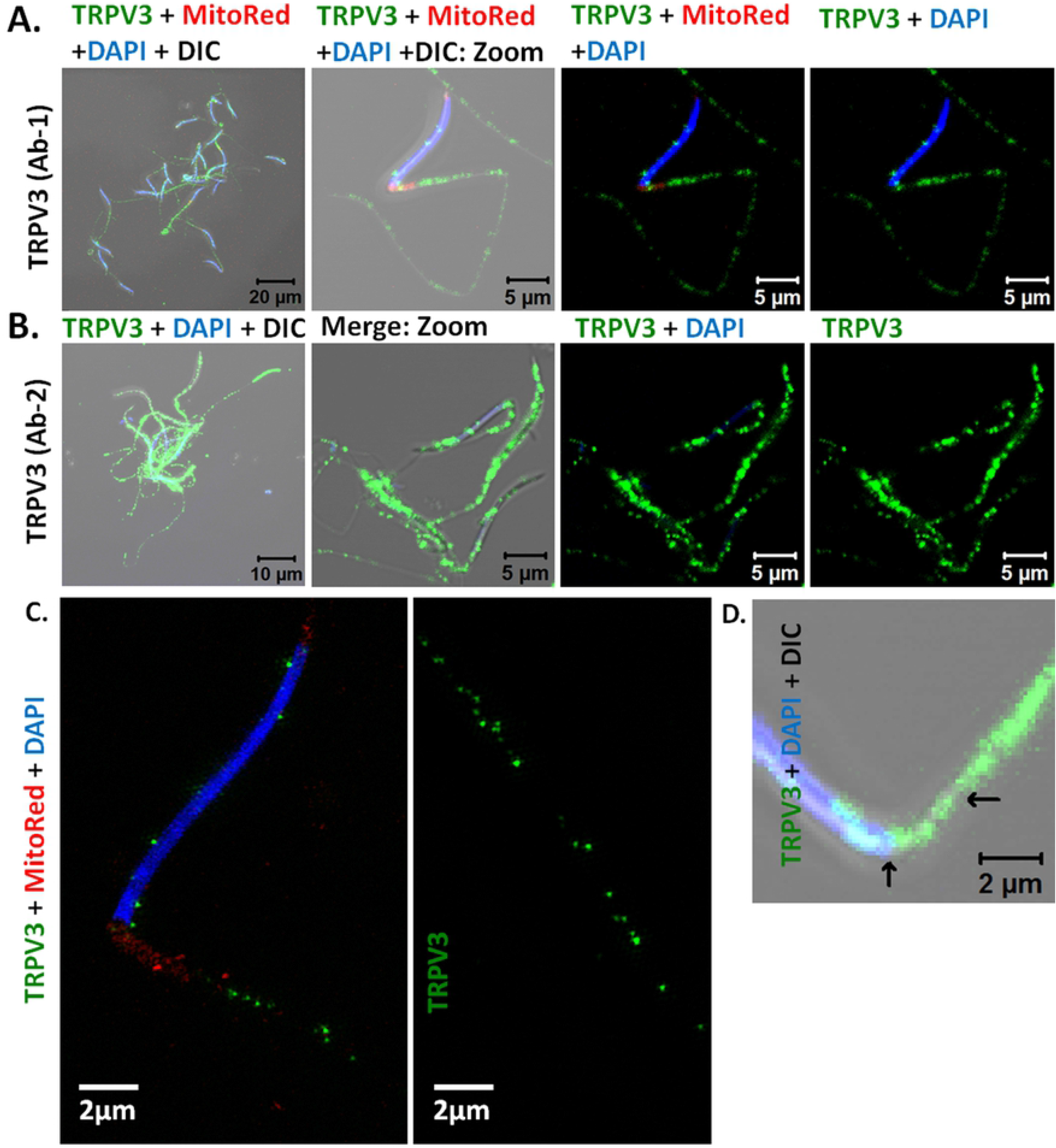
Microscopic images showing localization of TRPV3 in duck sperm. **A-B.** Confocal microscopic images depicting the localization of TRPV3 (green) as detected by two different antibodies and Nucleus (blue) by DAPI. Mitochondria (red) is labelled by Mitotracker Red dye in **A** and **C** to highlight the channel expression in the mitochondrial region. **C**. SR-SIM images of TRPV3 localization (using Ab1 antibody) at the head (left) and tail (right) of duck sperm is shown. **D.** Zoomed up image of neck region of sperm depicting the presence of TRPV3 (green) in the neck region. The head (blue) and arrows mark the start- and end-point of mitochondrial region.

Confocal microscopy using two different antibodies against the C-terminus of TRPV4 we found that TRPV4 is very scarce in the head, prominent at the mitochondrial region (seen by its colocalization with MitoTracker Red, indicated by white arrows) and at the tail of duck sperm (**Fig. 6A, B, D**). SIM-based super resolution imaging (using Ab1) revealed punctate distribution of TRPV4 throughout the sperm (**Fig. 6C**).

**Fig. 6.**
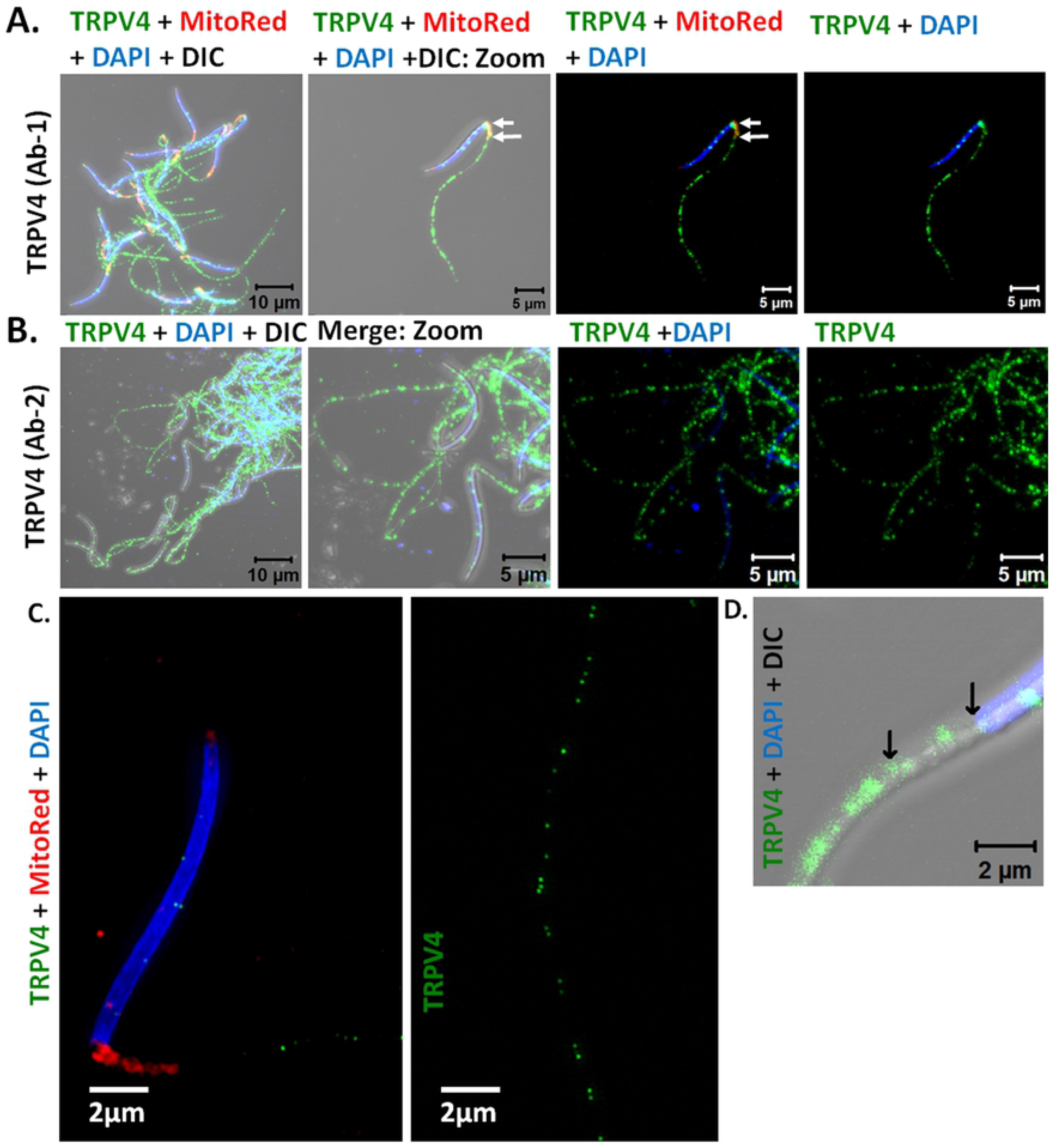
Microscopic images showing localization of TRPV4 in duck sperm. **(A-B)** Confocal microscopic images depicting the localization of TRPV4 (green) as detected by two different antibodies and Nucleus (blue) by DAPI. Mitochondria (red) is labelled by Mitotracker Red dye in **A** and **C** to highlight the channel expression in the mitochondrial region. (**C**) SR-SIM images of TRPV4 localization (using Ab1 antibody) at the head (left) and tail (right) of duck sperm is shown. (**D**) Zoomed up image of neck region of sperm depicting the presence of TRPV4 (green) in the neck region. The head (blue) and arrows mark the start and end point of mitochondrial region.

Confocal microscopy using two different antibodies against the C-terminus of TRPV5 we found that TRPV5 is exclusively present at the tail (indicated by white arrows), with highest density immediately after the mitochondrial region and intensity gradually decreasing towards the tapering end of the tail (**Fig. 7A, B, D**). TRPV5 is absent at the head and neck region. SIM-based super resolution imaging (using Ab1) revealed punctate distribution of TRPV5 at the tail of sperm (**Fig. 7C**).

**Fig. 7.**
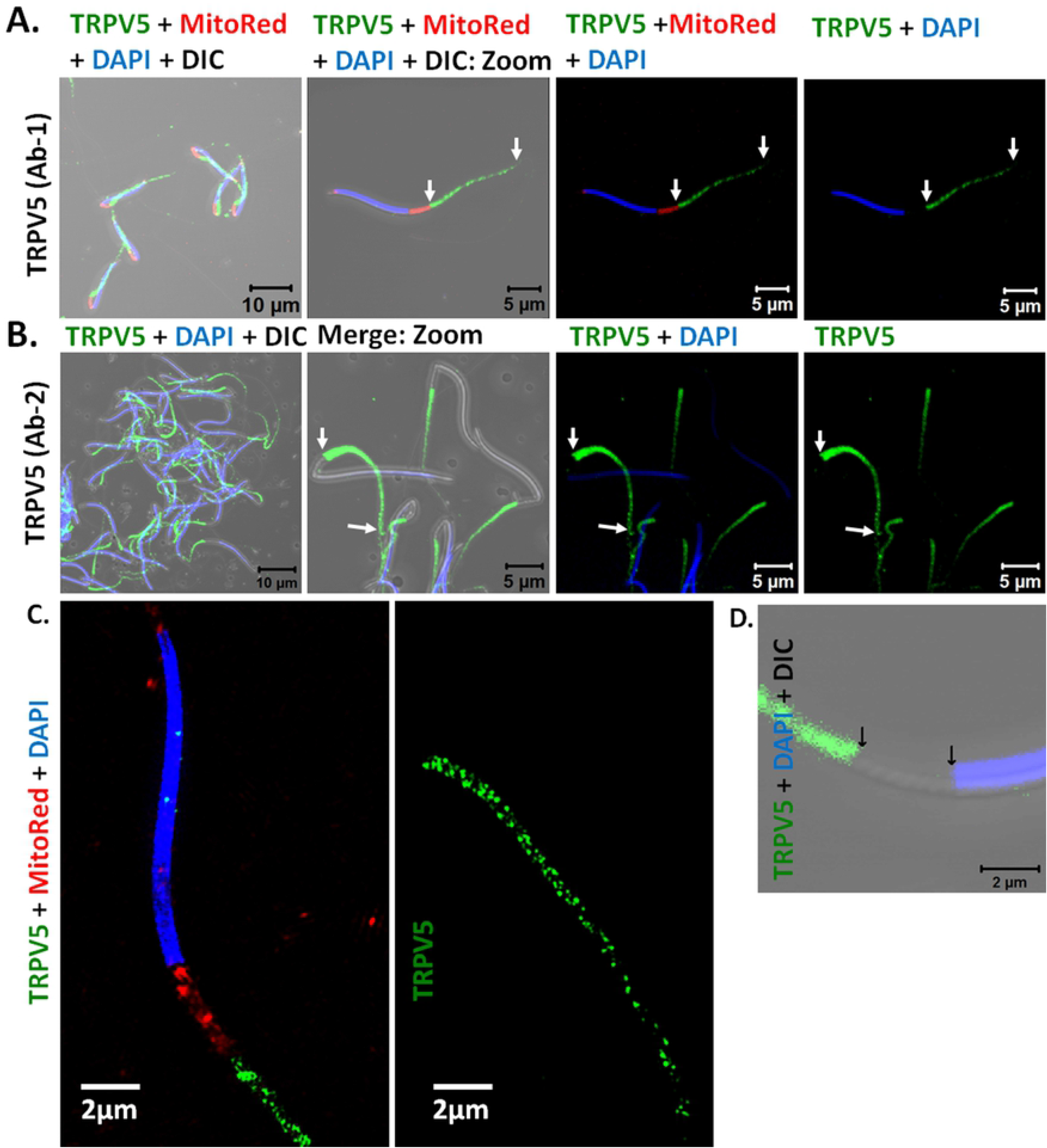
Microscopic images showing localization of TRPV5 in duck sperm. **(A-B)** Confocal microscopic images depicting the localization of TRPV5 (green) as detected by two different antibodies and Nucleus (blue) by DAPI. Mitochondria (red) is labelled by Mitotracker Red dye in **A** and **C** to highlight the channel expression in the mitochondrial region. (**C**) SR-SIM images of TRPV5 localization (using Ab1 antibody) at the head (left) and tail (right) of duck sperm is shown. (**D**) Zoomed up image of neck region of sperm depicting the absence of TRPV5 (green) in the neck region. The head (blue) and arrows mark the start and end point of mitochondrial region.

We detected the presence of TRPV6 throughout the sperm using confocal microscopy via two different antibodies against the C-terminus of TRPV6 (Ab1 and Ab2) (**Fig. 8A, B, D**). SIM-based super resolution imaging (using Ab1) revealed punctate distribution of TRPV6 throughout the sperm (**Fig. 8C**). Taken together, our results confirm the endogenous expression and differential localization of TRPVs in duck sperm.

**Fig. 8.**
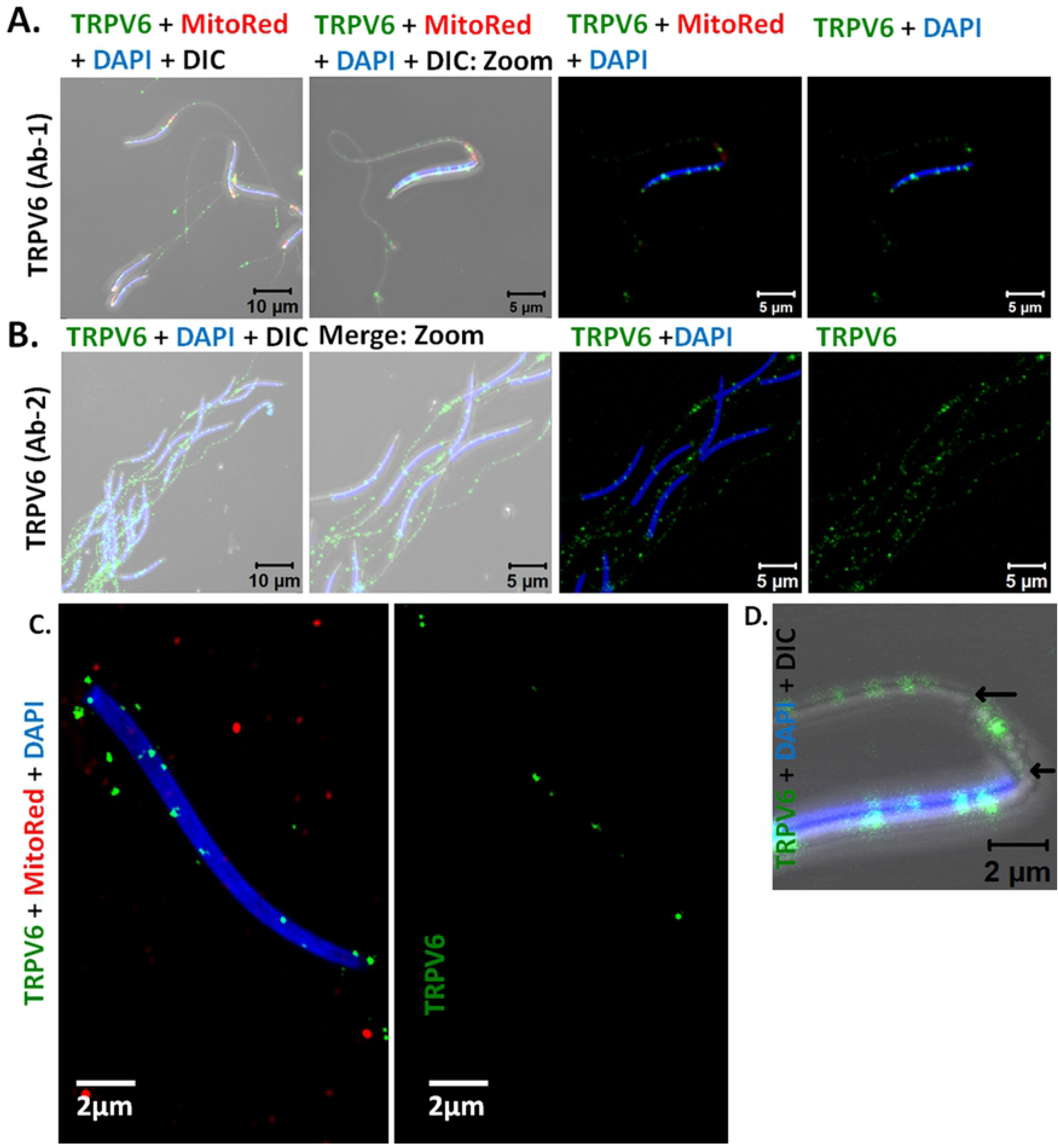
Microscopic images showing localization of TRPV6 in duck sperm. **(A-B)** Confocal microscopic images depicting the localization of TRPV6 (green) as detected by two different antibodies and Nucleus (blue) by DAPI. Mitochondria (red) is labelled by Mitotracker Red dye in **A** and **C** to highlight the channel expression in the mitochondrial region. (**C**) SR-SIM images of TRPV6 localization (using Ab1 antibody) at the head (left) and tail (right) of duck sperm is shown. (**D**) Zoomed up image of neck region of sperm depicting the absence of TRPV6 (green) in the neck region. The head (blue) and arrows mark the start and end point of mitochondrial region.

## 4. Discussion

Each part of the sperm is specialized to carry out specific functions during its movement in female reproductive tract and during fusion with the egg. The head contains DNA to be transferred to the egg, and the membrane of the head determine the optimal time for disintegrating to allow sperm-egg fusion [29]. Similarly the neck region contains mitochondria, which provides energy to the sperm, and membrane potential across this region is essential for proper mitochondrial functioning which affects sperm survivability and movement. The tail region is critical for generation of force, determination of speed and direction of sperm movement. Hence each of these sperm regions are likely to be equipped/enriched with specific ion channels that can sense physiological conditions like temperature, chemical cues, fluid pressure and accordingly conduct specific ion flux thus initiating signals in order to modulate appropriate response in the sperm. TRP channels form a unique group of ion channels that are multimodal in nature and respond to several physiological environment. Their presence at distinct location in sperm could determine their functional relevance in the fertilization process. Hence in this study we explored the presence and localization of TRPV channels, which includes four heat sensitive channels (TRPV1, TRPV2, TRPV3, TRPV4) and highly Ca^2+^-selective channels (TRPV5 and TRPV6) in duck sperm.

We have previously shown the presence of TRPV1 in the neck and tail region of fish sperm [16]. Pharmacological modulation of TRPV1 alters fish sperm motility and their ability to swim increases from 2 minutes to 90 minutes [16]. In this work, we have observed that TRPV1 is abundant at the neck and tail regions, indicating that TRPV1 could also regulate duck sperm motility. TRPV1 has also been shown to regulate thermotaxis of human sperm and this feature could be conserved in other warm blooded vertebrates like birds too [20]. TRPV1 has also been shown to regulate capacitation of other relevant species, at least for bull and boar sperm [18–19]. However, avian sperm are believed to not require any capacitation prior to fertilizing the egg [30–31]. Interestingly, TRPV1 activation mediated Ca^2+^-influx has been shown to mediate bovine sperm release from the oviductal epithelium [32]. If conserved in birds, this phenomenon could be critical in sustained release of fertilized eggs over a period of time post insemination, as the sperm gets stored and slowly released in the oviduct of birds [33]. Indeed our preliminary experiments suggest that TRPV1 activation may retain the motility of the bird sperm for a longer duration (**Supplementary movie 1**). These special oviductal structures, called sperm storage tubules (SSTs) is maintained at 41°C, a temperature higher than the body temperature of birds [33]. Notably, TRPV1 activation temperature in chickens is ~45°C, hence TRPV1 could be involved in oviductal release of duck sperm, a possibility that need to be verified in future [34].

So far TRPV2 and TRPV3 have not been reported in sperm of any species, hence their precise role in regulation of sperm functions is unknown. This work is the first report of the presence of TRPV2, TRPV3 in the sperm of any vertebrate. We demonstrate that TRPV2 and TRPV3 are also prominently present at the tail of duck sperm. Such localization may suggest the role of these two channels in the regulation of tail functions and in particular regulation of microtubule and actin cytoskeleton present there.

We reported the presence of TRPV4 in the sperm of vertebrates ranging from fishes to humans, and demonstrated that TRPV4 regulates Ca^2+^-levels in human sperm [24]. TRPV4 has been implicated in regulating murine sperm thermotaxis. Accordingly, the murine sperm can preferentially swim to higher temperatures (from 37°C to 38.4°C) in a TRPV4-dependent manner, as sperm from TRPV4^-/-^ animals (TRPV4 knock-out) didn’t show this thermo-sensitivity [22]. Recently, TRPV4 has been confirmed as a temperature-sensitive ion channel of human sperm, and TRPV4 activation has been demonstrated to trigger the initial membrane depolarization in sperm, facilitating CatSper and Hv1 activation and, subsequently resulting in sperm hyperactivation, a phenomenon essential for sperm-egg fusion [23]. Based on these rent work, it is possible that in duck too, TRPV4 can regulate critical aspects of sperm physiology and can have a major impact on fertility in duck.

TRPV5 and TRPV6 channels are not thermosensitive, but are highly Ca^2+^ selective ion channels. So far TRPV5 hasn’t been reported in the sperm of any other species, except rat. In rat, TRPV5 express in the spermatogenic cells and in spermatozoa [35]. In this work, we report that TRPV5 has the highest intensity among all TRPV channels. TRPV5 expression is also very specific, it starts with highest density immediately after the mitochondrial region and gradually decreases towards the tapering end of the tail. Such prominent expression of TRPV5 in duck sperm tail can be strongly correlated with the motility functions performed by the tail region.

Similarly, we demonstrate that TRPV6 is also present in the duck sperm. TRPV6 is localized throughout the duck sperm, however among all TRPV channels tested in this work, the expression of TRPV6 is quite low. Recent reports suggest that mutations rendering inactivation of TRPV6 or excision of the pore-forming region and the complete cytosolic C terminus of TRPV6 (thus rendering it non-functional) leads to infertility in male mice [36–37]. Both these studies have reinforced that TRPV6-mediated decrease in the extracellular Ca^2+^ concentration near the distal epididymal duct is essential for the acquisition of physiological functions and survival of mice spermatozoa. If the role of TRPV6 remain same in duck sperm too, then TRPV6 could be a major factor controlling duck sperm fertilization ability. Taken together, we demonstrate the endogenous expression, distinct localization and compartmentalization of thermosensitive and Ca^2+^-selective TRPV channels in mature duck sperm. The endogenous presence and distinct distributions can be correlated to the known functions of these channels in general and also specific functions reported in the sperm of other vertebrates.

Sperm cell proteome is very different than most of the other tissues as testis, spermatozoa and mature sperm contain many proteins that are not expressed in any other tissues/cells. Many of these proteins are actually due to alternate splicing and/or post translational modifications. The complexity of sperm proteome is still poorly understood [38–39]. Understanding of such proteome and changes in such proteomes have huge importance in the context of fertility and sterility [40]. In this work, we demonstrate that at least one detectable form of TRPV1-TRPV6 is expressed in mature duck sperm. This understanding may also have impact on the commercialization of the poultry birds and conservation of rare-endangered species globally.

## Author contributions and notes

SCG collected the mature sperm from duck. RKM and AK did all the experiments related to western blot, antibody staining, flow-cytometry and confocal imaging. CG did all the SIM-based imaging. RKM and CG wrote the manuscript.

## Acknowledgements

This work was supported by intramural funding to NISER by Department of Atomic Energy, Govt. of India. We are thankful to the Imaging and Flow cytometry facility of NISER. We are thankful to Carl-Zeiss India, Bangalore for their help in performing super-resolution imaging using ELYRA SR-SIM.

## List of abbreviations

BSA: Bovine Serum Albumin
DAPI: 4′,6-diamidino-2-phenylindole
DIC: Differential Interference Contrast
DNA: Deoxyribo nucleic acid
FACS buffer: Flow cytometry buffer
MFI: Mean Fluorescence Intensity
MitoRed: Mitotracker red
NaCl: Sodium Chloride
PBS: Phosphate buffer saline
PBS-T: 0.1% Tween 20 in PBS
PFA: Paraformaldehyde
PVDF: Polyvinylidene fluoride
SDS-PAGE: Sodium Dodecyl Sulfate Poly Acrylamide Gel Electrophoresis
SR-SIM: Super Resolution Structured Illumination Microscopy
TRP: Transient Receptor Potential
TRPV: Transient Receptor Potential Vanilloid
WB: Western Blot analysis

## References

1. Yoshida M., Kawano N., Yoshida K. Control of sperm motility and fertility: diverse factors and common mechanisms. Cell Mol Life Sci. 2008; 65: 3446–3457.

2. Bahat A., Tur-Kaspa I., Gakamsky A., Giojalas L.C., Breitbart H., Eisenbach M. Thermotaxis of mammalian sperm cells: a potential navigation mechanism in the female genital tract. Nat Med. 2003; 9: 149–150.

3. Saha S., Sucharita S., Majhi R.K., Tiwari A., Pradhan S.K., Patra B.K., et al. TRPA1 is selected as a semi-conserved channel during vertebrate evolution due to its involvement in spermatogenesis. Biochem Biophys Res Commun. 2019; 512: 295–302.

4. Bahat A., Caplan S.R., Eisenbach M. Thermotaxis of human sperm cells in extraordinarily shallow temperature gradients over a wide range. PLoS One. 2012; 7: e41915.

5. Pérez-Cerezales S., Laguna-Barraza R., de Castro A.C., Sánchez-Calabuig M.J., Cano-Oliva E., de Castro-Pita F.J., et al. Sperm selection by thermotaxis improves ICSI outcome in mice. Sci Rep. 2018; 8: 2902.

6. Grunewald S., Paasch U., Glander H.J. & Anderegg U. Mature human spermatozoa do not transcribe novel RNA. Andrologia 2005; 37: 69–71.

7. Asano A., Nelson J.L., Zhang S. & Travis A.J. Characterization of the proteomes associating with three distinct membrane raft sub-types in murine sperm. Proteomics 2010; 10: 3494–3505.

8. Goodrich R.J., Anton E. & Krawetz S.A. Isolating mRNA and small noncoding RNAs from human sperm. Methods Mol. Biol. 2013; 927: 385–396.

9. Eisenbach M. Sperm chemotaxis. Rev. Reprod. 1999; 4: 56–66.

10. Spehr M., Gisselmann G., Poplawski A., Riffell J.A., Wetzel CH, Zimmer RK & Hatt H. Identification of a testicular odorant receptor mediating human sperm chemotaxis. Science 2003; 299: 2054–2058.

11. Carlson A.E., Westenbroek R.E., Quill T., Ren D., Clapham D.E., Hille B., et al. CatSper1 required for evoked Ca2+ entry and control of flagellar function in sperm. Proc. Natl. Acad. Sci. U. S. A. 2003; 100: 14864–14868.

12. Suarez S.S. & Ho H.C. Hyperactivated motility in sperm. Reprod. Domest. Anim. 2003; 38: 119–124.

13. Kirkman-Brown J.C., Punt E.L., Barratt C.L.R. & Publicover S.J. Zona pellucida and progesterone-induced Ca2+ signaling and acrosome reaction in human spermatozoa. J. Androl. 2002; 23: 306–315.

14. Nilius B., Owsianik G. The transient receptor potential family of ion channels. Genome Biol. 2011; 12: 218.

15. Wu L.J., Sweet T.B., Clapham D.E. International Union of Basic and Clinical Pharmacology. LXXVI. Current progress in the mammalian TRP ion channel family. Pharmacol Rev. 2010; 62: 381–404.

16. Majhi R.K., Kumar A., Yadav M., Swain N., Kumari S., Saha A. Thermosensitive ion channel TRPV1 is endogenously expressed in the sperm of a fresh water teleost fish (*Labeo rohita*) and regulates sperm motility. Channels (Austin). 2013; 7: 483–492.

17. Grimaldi P., Orlando P., Di Siena S., Lolicato F., Petrosino S., Bisogno T., et al. The endocannabinoid system and pivotal role of the CB2 receptor in mouse spermatogenesis. Proc Natl Acad Sci U S A. 2009; 106: 11131–11136.

18. Maccarrone M., Barboni B., Paradisi A., Bernabò N., Gasperi V., Pistilli M.G., et al. Characterization of the endocannabinoid system in boar spermatozoa and implications for sperm capacitation and acrosome reaction. J Cell Sci. 2005; 118: 4393–4404.

19. Gervasi M.G., Osycka-Salut C., Caballero J., Vazquez-Levin M., Pereyra E., Billi S., et al. Anandamide capacitates bull spermatozoa through CB1 and TRPV1 activation. PLoS One. 2011; 6: e16993.

20. De Toni L., Garolla A., Menegazzo M., Magagna S., Di Nisio A., Šabović I., et al. Heat Sensing Receptor TRPV1 Is a Mediator of Thermotaxis in Human Spermatozoa. PLoS One. 2016; 11: e0167622.

21. Francavilla F., Battista N., Barbonetti A., Vassallo M.R., Rapino C., Antonangelo C., et al. Characterization of the endocannabinoid system in human spermatozoa and involvement of transient receptor potential vanilloid 1 receptor in their fertilizing ability. Endocrinology 2009; 150: 4692–4700.

22. Hamano K., Kawanishi T., Mizuno A., Suzuki M., Takagi Y. Involvement of Transient Receptor Potential Vanilloid (TRPV) 4 in mouse sperm thermotaxis. J Reprod Dev. 2016; 62: 415–422.

23. Mundt N., Spehr M., Lishko P.V. TRPV4 is the temperature-sensitive ion channel of human sperm. Elife. 2018; 7: pii: e35853. doi: 10.7554/eLife.35853.

24. Kumar A., Majhi RK, Swain N., Giri S.C., Kar S., Samanta L., et al. TRPV4 is endogenously expressed in vertebrate spermatozoa and regulates intracellular calcium in human sperm. Biochem Biophys Res Commun. 2016; 473: 781–788.

25. Majhi R.K, Kumar A., Yadav M., Kumar P., Maity A., Giri S.C., et al. Light and electron microscopic study of mature spermatozoa from White Pekin duck (*Anas platyrhynchos*): an ultrastructural and molecular analysis. Andrology. 2016; 4: 232–244.

26. Majhi R.K., Saha S., Kumar A., Ghosh A., Swain N., Goswami L., et al. Expression of temperature-sensitive ion channel TRPM8 in sperm cells correlates with vertebrate evolution. PeerJ. 2015; 3: e1310.

27. Laemmli U.K. Cleavage of structural proteins during the assembly of the head of bacteriophage T4. Nature 1970; 227: 680–685.

28. Somogyi C.S., Matta C., Foldvari Z., Juhász T., Katona É., Takács Á.R., Hajdú T., Dobrosi N., Gergely P., Zákány R. Polymodal Transient Receptor Potential Vanilloid (TRPV) Ion Channels in Chondrogenic Cells. Int J Mol Sci. 2015; 16: 18412–18438.

29. Inoue N., Hamada D., Kamikubo H., Hirata K., Kataoka M., Yamamoto M., et al. Molecular dissection of IZUMO1, a sperm protein essential for sperm-egg fusion. Development. 2013; 140: 3221–3229.

30. Howarth B Jr, An examination for sperm capacitation in the fowl. Biol Reprod. 1970; 3: 338–341.

31. Nixon B., Ewen K.A., Krivanek K.M., Clulow J., Kidd G., Ecroyd H., Jones R.C. Post-testicular sperm maturation and identification of an epididymal protein in the Japanese quail (*Coturnix coturnix japonica*). Reproduction. 2014; 147: 265–277.

32. Gervasi M.G., Osycka-Salut C., Sanchez T., Alonso C.A., Llados C., Castellano L., et al. Sperm Release From the Oviductal Epithelium Depends on Ca(2+) Influx Upon Activation of CB1 and TRPV1 by Anandamide. J Cell Biochem. 2016; 117: 320–333.

33. Sasanami T., Matsuzaki M., Mizushima S., Hiyama G. Sperm storage in the female reproductive tract in birds. J Reprod Dev. 2013; 59, 334–338.

34. Jordt S.E., Julius D. Molecular basis for species-specific sensitivity to “hot” chili peppers. Cell. 2002; 108: 421–430.

35. Li S., Wang X., Ye H., Gao W., Pu X., Yang Z. Distribution profiles of transient receptor potential melastatin- and vanilloid-related channels in rat spermatogenic cells and sperm. Mol Biol Rep. 2010; 37: 1287–1293.

36. Weissgerber P., Kriebs U., Tsvilovskyy V., Olausson J., Kretz O., Stoerger C. et al. Male fertility depends on Ca^2+^ absorption by TRPV6 in epididymal epithelia. Sci Signal. 2011; 4: ra27.

37. Weissgerber P., Kriebs U., Tsvilovskyy V., Olausson J., Kretz O., Stoerger C. et al. Excision of Trpv6 gene leads to severe defects in epididymal Ca^2+^ absorption and male fertility much like single D541A pore mutation. J Biol Chem. 2012; 287: 17930–17941.

38. Gilany K., Minai-Tehrani A., Amini M., Agharezaee N., Arjmand B. The Challenge of Human Spermatozoa Proteome: A Systematic Review. J Reprod Infertil. 2017; 18, 267–279.

39. Aitken R.J., Baker M.A. The role of proteomics in understanding sperm cell biology. Int J Androl. 2008; 31, 295–302.

40. Li C.J., Wang D., Zhou X. Sperm proteome and reproductive technologies in mammals. Anim Reprod Sci. 2016; 173: 1–7.

